# Passage through micro-sprayer increases functional activity – implications for activity assays in time-resolved cryo-EM

**DOI:** 10.1101/2024.12.01.624867

**Authors:** Priyanka Garg, Xiangsong Feng, Swastik De, Joachim Frank

## Abstract

This study examines the validity of an assay that is used to report on the retainment of functional competence by ribosomes as they pass a micro-sprayer. We find a reproducible *increase*, rather than the expected *decrease* in GFP production as monitored by fluorescence, which suggests heterogeneity or partial aggregation of ribosomes in solution. An even larger increase in functional activity is observed when sonication is used, pointing to mechanical agitation as the decisive factor in both scenarios. The results have a bearing on the design and interpretation of validation experiments in time-resolved cryo-EM based on microfluidic chips.

## Introduction

Single-particle cryo-electron microscopy (cryo-EM) has revolutionized structural biology by allowing scientists to visualize biological macromolecules in their native states at high resolutions. Following a widely practised protocol, a thin layer of liquid sample is applied to the EM grid by pipetting and blotting, after which the grid is rapidly immersed in a cryogen, resulting in molecules embedded in vitreous ice (Adrian et al., 1984; Armstrong et al., 2020; Grassucci et al., 2007; Kaledhonkar et al., 2018; Kühlbrandt, 2014; Snijder et al., 2017). An alternative sample preparation technique involves spraying the sample onto the grid, allowing droplets to spread to the right thickness (Ashtiani et al., 2018; Avvaru et al., 2006; Berriman and Unwin, 1994; Feng et al., 2017; Klebl et al., 2020; Kontziampasis et al., 2019; Rubinstein et al., 2019; White et al., 2003; Yang et al., 2024). This alternative technique may have advantages over the traditional method as it shortens the time during which the molecules are exposed to the air-water interface (Klebl et al., 2020; Yang et al., 2024). Importantly, spraying as a method of sample deposition is used in several experimental approaches to time-resolved single-particle cryo-EM where a biomolecular reaction is realized in a microfluidic chip, the reaction product is sprayed onto the EM grid, and the grid is plunged into the cryogen (Lu et al., 2008). Different time points are sampled by stopping the reaction at different times, in the range of milliseconds, offering the opportunity to visualize intermediate states.

Our study was motivated by concerns about the damage to biological molecules inflicted in the experiments. In their passage through the sprayer, biological molecules (proteins, RNAs, and ribonucleoprotein complexes [RNPs]) are subject to strong forces (such as sheering and torque forces), and this raises the question of whether they retain their structural and functional integrity in this mode of sample preparation. Thus the validity of results obtained in any experiments utilizing spray deposition time-resolved or otherwise is at stake.

In the approach to time-resolved cryo-EM we have adopted (Lu et al., 2008), early validation experiments were done in the Agrawal lab (Shaikh et al., 2014) with *E. coli* ribosomes. In that study, a cell-free translation assay was performed on the spray mixture collected, to assess whether passage through the microfluidic device preserved ribosomal translation activity. Intact 70S ribosomes were sprayed through the device at three different concentrations, and their translation activity, measured by fluorescence of newly synthesised GFP, was compared to that of ribosomes that had not passed through the device. Their findings indicated that passing pre-associated 70S ribosomes formed by incubating purified 30S and 50S subunits through the microfluidic device did not impair translation activity. While two out of three runs of the experiment showed only slight changes in activity, one demonstrated an enhancement by 20%. This unexpected enhancement, dismissed by these authors as an outlier, foreshadows the results of our present study.

As we recently replaced silicon with PDMS in the fabrication of microfluidic chips (Bhattacharjee et al., 2024), there was a need to revisit the question of validity, prompting us to repeat the ribosome activity test of the Agrawal lab. Our tests showed a consistent, highly significant increase in the functional activity of ribosomes after passage through the sprayer. Moreover, an even larger increase was observed following the sonication of the sample, indicating that different forms of mechanical agitation render the ribosomes in the sample more active. This result has implications for the validation of experiments in several setups which use piezo-based sonic vibration for sample deposition (Jain et al., 2012; Rubinstein et al., 2019).

## Methods

### Preparation of template

The Neon green sequence-containing vector, which served as the template for PCR, was synthesised by Integrated DNA Technologies (Coralville, Iowa). The reaction was conducted using Forward (GCCCCGCGAAATTAATACG) and Reverse (CCTCCTTTCAGCAAAAAACC) primers with an annealing temperature of 50°C. The resulting amplified linear product was subsequently utilized as the template.

### Preparation of the sample using the TR micro-sprayer

We employed an experimental setup comprising an environmental chamber, a micro-sprayer, and a sample injection unit as used by Bhattacharjee *et. al*. (Bhattacharjee et al., 2024). The temperature and humidity were maintained at 22-24°C and 90-95%, respectively. *E. coli* 70S ribosome, comprising the large 50S and small 30S subunits, was obtained from New England Biolabs (NEB) (catalogue number P0763S) with an initial concentration of 13.3 µM. The sample was diluted to 7.2 µM using a buffer solution containing 20 mM Tris-HCl (pH 7.5), 100 mM NH4Cl, 10 mM Mg(OAc)2, and 4 mM βME. The syringe and tubing of the sample injection unit were first washed with 3*50µl of filtered water, followed by equilibration with the same buffer solution used for protein dilution. 25 µl of ribosome sample (7.2 µM) was then loaded into a glass syringe, which was further diluted to a concentration of 4-4.5 µM to account for the dead volume of the valve unit (25-30µl). A software-controlled system was used to control the flow rates (Fig. 1A).

**Figure 1:**
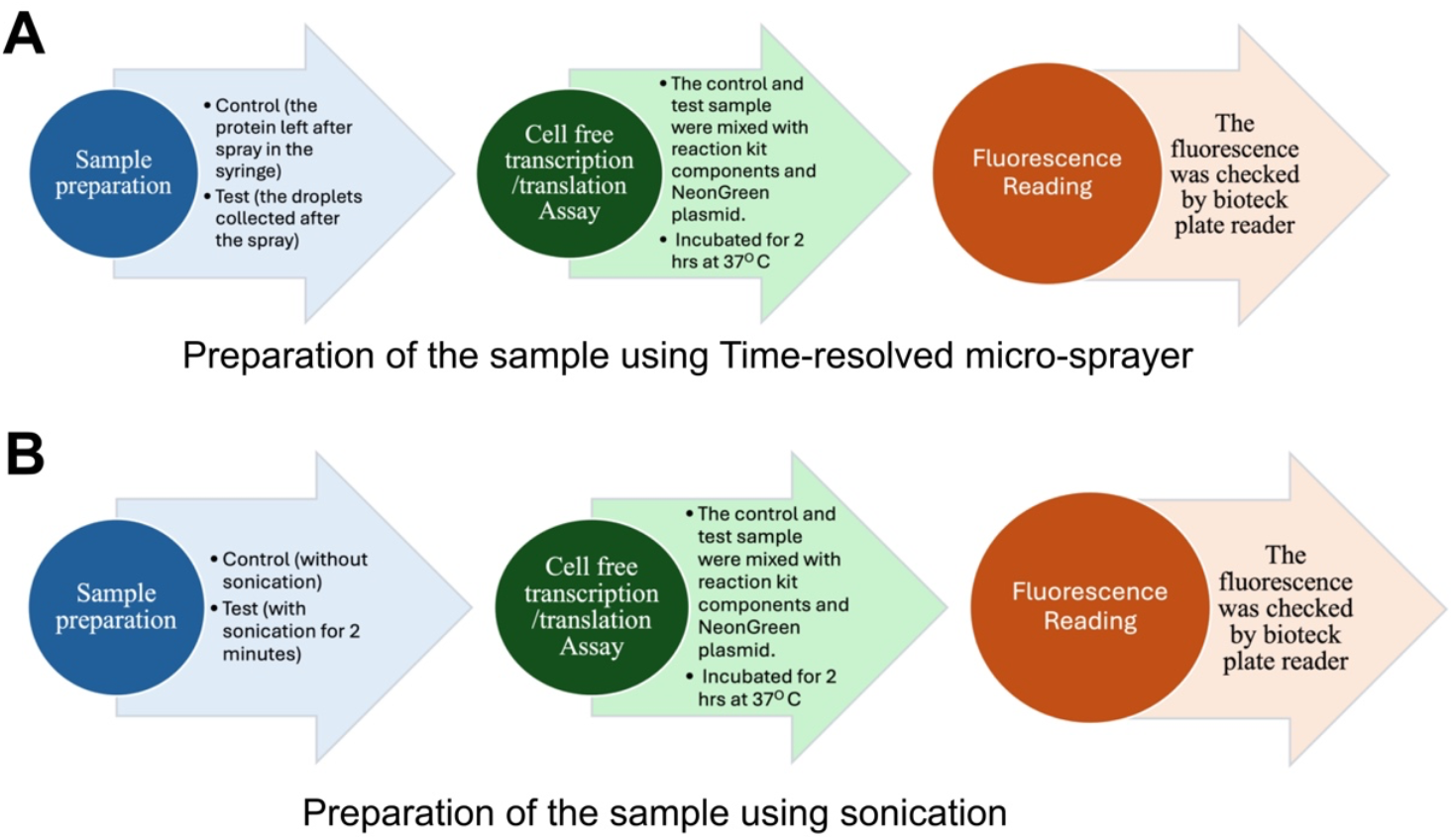
Steps of checking the activity of ribosomes. (A) before and after passage through the micro-sprayer; (B) before and after sonication.

### Preparation of the sample using sonication

The *E. coli* 70S ribosome sample with an initial concentration of 13.3 µM was diluted to 4.2 µM. Then the sample was sonicated at 25°C for 2 minutes by using Branson1510R-MT Ultrasonic Cleaner at a frequency of 40 kHz. Ribosome activity before and after sonication was compared (Fig. 1B).

## Results

### Experimental setup

The micro-sprayer was positioned inside the environmental chamber, with its nozzle directed toward an Eppendorf tube for sample collection (Fig. 2). The ribosome sample was sprayed through the nozzle under two different pressure conditions: 2 psi and 8 psi, with a flow rate of 5 µl/s, thereby ensuring that no dripping occurred (Feng et al., 2017). The ribosome concentration was measured before and after spraying.

**Figure 2:**
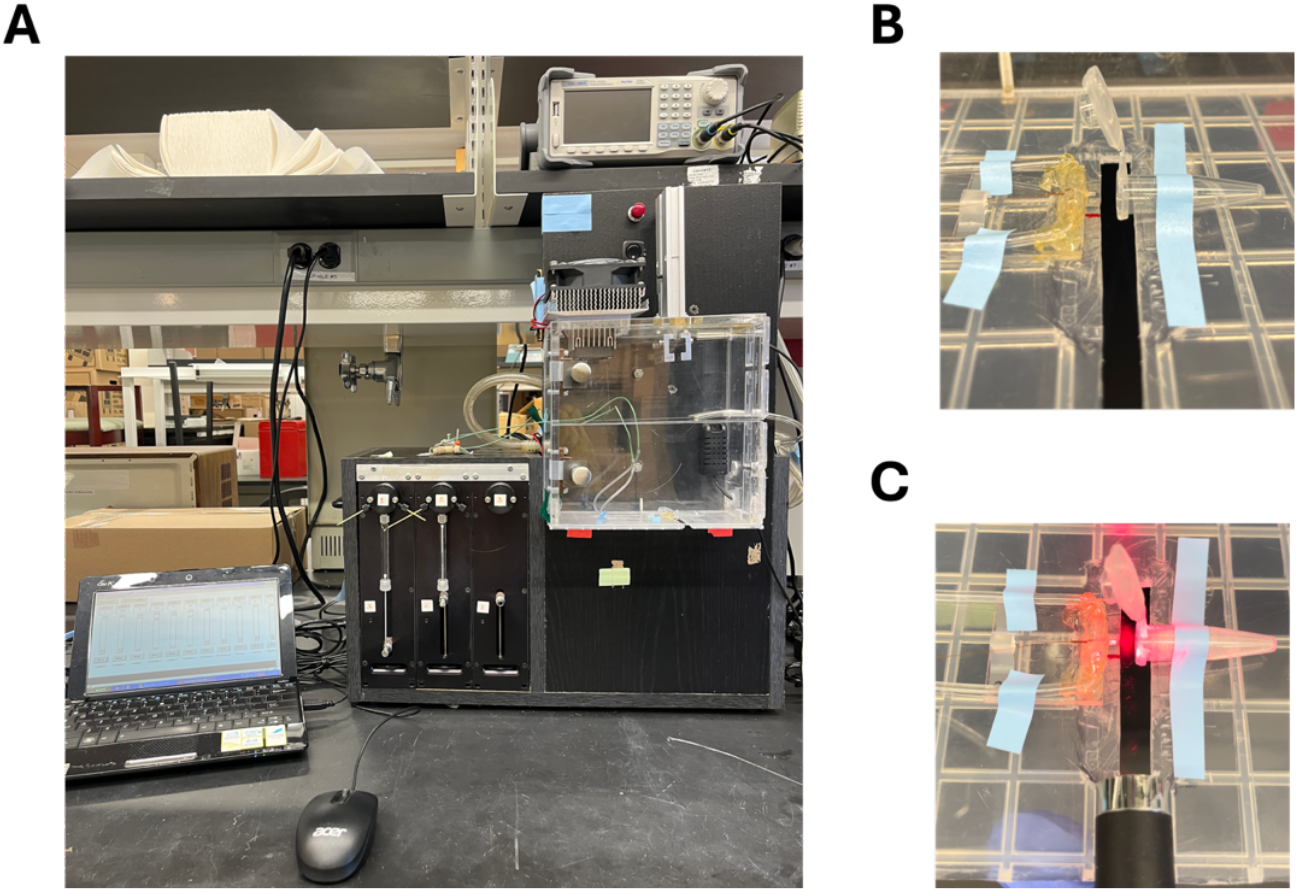
Experimental setup:(A) Apparatus of time-resolved cryo-EM sample preparation; (B) Enlarged view before spray; (C) Enlarged view of sample collection on the wall of an Eppendorf tube.

### Activity assay

We assessed the activity of the 70S *E. coli* ribosomes using a cell-free translation system (PURExpress^®^ Δ Ribosome Kit, New England Biolabs (NEB), Ipswich, Massachusetts). Samples collected before and after spraying, as well as before and after sonication, and negative control without ribosomes were mixed with solution A and buffer B from the kit, along with Neon green linear DNA, to a final protein concentration of 2.4 µM and a total volume of 12.5 µl. The mixtures were incubated at 37°C for 2 hours. Before fluorescence measurement, the volume of each sample was adjusted to 100 µl with additional buffer. Fluorescence readings were taken at excitation and emission wavelengths of 490 and 517 nm, respectively, using a 96-well plate. In addition to the experimental samples, we included a negative control to validate our findings. The negative control consisted of a sample buffer devoid of ribosomes, which was subjected to the same experimental procedure.

### Effect of spraying on the activity of the ribosome

We measured the fluorescence before and after spraying the sample under two different pressure conditions, 2 and 8 psi. Remarkably, after performing the experiment 3 times, our results (Fig. 3) showed a significant reproducible increase of 12 % and 15 % in sample “After, 2psi” and sample “After, 8psi” in ribosome activity, respectively, as evidenced by a higher fluorescence value compared to measurements taken before spraying. Notably, ribosome activity increased slightly more (but within the error margin) at the lower pressure of 2 psi than at 8 psi. The negative control confirmed that the observed fluorescence increase was due to the increase in activity of the ribosomes and was not attributable to any change in background fluorescence or the presence of experimental artefacts.

**Figure 3:**
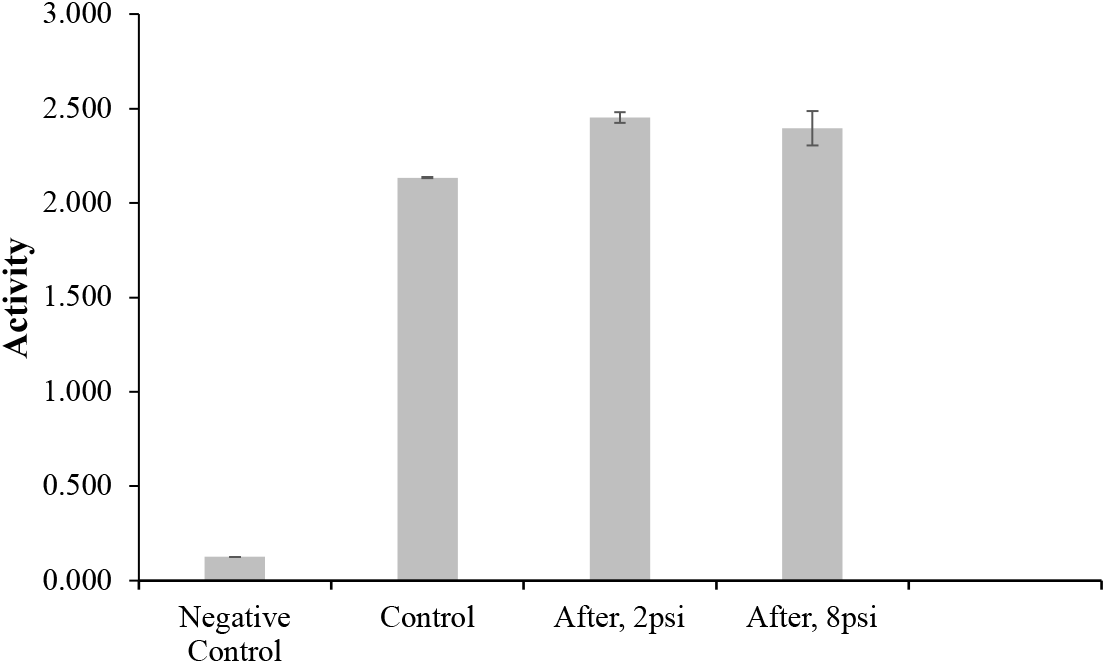
Effect of spraying on the activity of the ribosome. Negative control sample was without ribosomes; control sample was collected with ribosomes before spraying; samples labeled ‘After,2psi’ and ‘After,8psi’ were collected with ribosomes after spraying with ∼2 psi and 8 psi pressure respectively.

### Effect of sonication on the activity of the ribosome

The reproducible increase of functional activity of the ribosome after passage through the sprayer led us to hypothesize that mechanical agitation of any kind might have the same effect. To test this hypothesis we subjected the ribosome sample to sonication and performed the same assay, measuring GFP fluorescence before and after the treatment. Our results (Fig. 4) demonstrated again a substantial increase in ribosome activity following sonication, but this time it was more than double (by 37% vs 15%) of what was observed with the sprayer.

**Figure 4:**
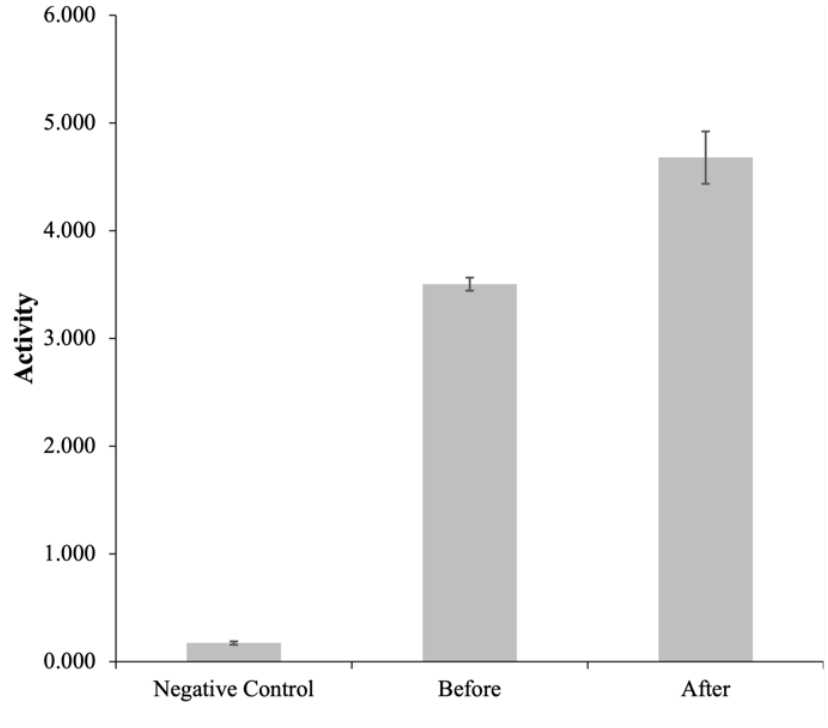
Effect of sonication on the activity of ribosome. Negative control sample is without ribosome; samples labeled “Before” and “After” were collected before and after sonication, respectively.

## Discussion

The spraying process utilized in time-resolved cryo-EM offers significant advantages over traditional blotting techniques, particularly in mitigating the detrimental effects of the air-water interface on sample quality. In conventional blotting methods, samples are often exposed to this interface during air drying, leading to issues such as protein denaturation and aggregation. In contrast, the spraying process minimizes this exposure, allowing much quicker and more efficient sample deposition. This reduction is critical for maintaining protein integrity and functionality, which are essential for accurate structural analysis.

However, the strong mechanical forces molecules are subjected to during spraying raise concerns about artefacts that might be introduced in this process. In experiments where spraying is solely used for the deposition of a sample, as an alternative to pipetting/blotting, such artefacts may have to be considered in comparing results obtained from either method. In time-resolved experiments using a micro-sprayer, the validity of multiple observations of intermediate states is at stake.

With a molecule as complex as the ribosome, we expected a *decrease* in functional activity, as measured by fluorescence of GFP newly synthesized on the ribosome either before or after passage through the spray nozzle. What we found instead was a significant *increase*. Such an increase in activity was in fact reported in one of three experiments of a similar kind by the Agrawal group (Shaikh et al., 2014), but it was dismissed as an outlier. We hypothesized that the increase might be the result of mechanical agitation during passage and that sonication would give a similar effect. A straightforward experiment confirmed that this was indeed the case, even with more than twice the increase observed with the sprayer (37% vs 15%).

As a possible explanation, mechanical agitation may facilitate favourable conformational changes, leading to more efficient substrate binding and catalytic activity, or in some other way changing the balance between pre-existing active and inactive ribosome populations. The forces produced by spraying or sonication might also be effective in disrupting aggregations of ribosomes, restoring them to their active conformations as this would increase the availability of active sites (Jain et al., 2012; Rubinstein et al., 2019). This alternative explanation is suggested by the fact that sonication has been often shown to enhance protein activity by disrupting aggregates and improving mixing, which in turn increases the availability of active sites (Ashokkumar et al., 2009; Zhu et al., 2018).

As a take-home message, the results of functional activity assays, which are essential for the validation of new technology such as time-resolved cryo-EM, need to be viewed with some caution since they make assumptions about the homogeneity of the sample in terms of functional competence and aggregation state that may not always be fulfilled.

## Acknowledgements

This work was supported by a grant from the National Institutes of Health (R35GM139453) to J.F. I thank Sanjay Saini for his assistance with the fluorescence lab instrument and Riley Charles Gentry for sharing valuable insights regarding the interpretation of our results.

## Notes

### Competing Interest Statement

The authors have declared no competing interest.

